# DIAPH3 deficiency links microtubules to mitotic errors, defective neurogenesis, and brain dysfunction

**DOI:** 10.1101/2020.08.11.245829

**Authors:** Eva On-Chai Lau, Devid Damiani, Yves Jossin, Georges Chehade, Olivier Schakman, Nicolas Tajeddine, Philippe Gailly, Fadel Tissir

**Author notes:** Correspondence to: Fadel TISSIR, Avenue Mounier 73 Box B1.73.16, 1200 Brussels Belgium.

## Abstract

Diaphanous (DIAPH) 3 is a member of the formin proteins that have the capacity to nucleate and elongate actin filaments and therefore, to remodel the cytoskeleton. DIAPH3 is essential for cytokinesis as its dysfunction impairs the contractile ring and produces multinucleated cells. Here, we report that DIAPH3 localizes at the centrosome during mitosis and regulates the assembly and polarity of the mitotic spindle. DIAPH3-deficient cells display disorganized cytoskeleton, multipolar spindles, and supernumerary centrosomes. DIAPH3-deficiency disrupts the expression and/or stability of microtubule-associated proteins SPAG5 and KNSTRN. SPAG5 and DIAPH3 have similar expression patterns in the developing brain and overlapping subcellular localization during mitosis. Knockdown of SPAG5 phenocopies the DIAPH3 deficiency, whereas its overexpression rescues the DIAH3 phenotype. Conditional inactivation of *Diaph3* in the cerebral cortex profoundly disrupts neurogenesis depleting cortical progenitors and neurons; and leading to cortical malformation and autistic-like behavior. Our data uncover uncharacterized functions of DIAPH3 and provide evidence that this protein belongs to a molecular toolbox that links microtubule dynamics during mitosis to aneuploidy, cell death, fate determination defects, and cortical malformation.

## Introduction

Development of the cerebral cortex requires the production and positioning of the right number of neurons. At initial stages of cortical development, the dorsal telencephalon is organized in a pseudostratified epithelium consisting of neural stem cells (NSCs, also known as neuroepithelial cells) that undergo multiple rounds of proliferative/symmetric division to expand the initial pool of progenitors. Once neurogenesis begins, the neocortex comprises two germinal zones: the ventricular zone (VZ), which forms the lining of lateral ventricles and contains radial glial cells (RG), also known as apical neural progenitor cells (aNPC); and the adjacent subventricular zone (SVZ), which is located dorsally to the VZ and contains basal progenitors (BP). In VZ, aNPC undergo several rounds of divisions to self-renew and generate glutamatergic neurons. aNPC cells can also give rise to BP cells, which delaminate from the VZ and translocate to SVZ where they divide a limited number of times to increase the final output of neurons (Florio and Huttner, 2014, Noctor et al., 2008). A delicate balance between proliferation and differentiation of aNPC must be preserved during neurogenesis. This balance is regulated by intrinsic and extrinsic factors, and involves rearrangements of the cytoskeleton to support a rigorous sequence of fate decisions. During cell division, filamentous actin rearranges at the cell cortex to enhance cell membrane rigidity for anchorage of astral microtubules (Heng and Koh, 2010). In the meantime, the centrosome, also known as the microtubule organising centre (MTOC), duplicates and the two nascent centrosomes migrate toward the poles of the cell. Astral and spindle microtubules nucleate from the centrosomes and extend to cortex and equator of the cell respectively. Polarity proteins GPSM2 (aka PINS/LGN) and NUMA are distributed underneath the cell cortex and interact with cortical actin to connect microtubule plus-end motor proteins dynein/dynactin, and pull on astral microtubules (Morin and Bellaiche, 2011). On the other hand, spindle microtubules grow inwardly and attach to chromosomes at the metaphase plate. Once the chromosomes are properly aligned and each chromosome is bilaterally connected to two spindle microtubules, cohesin is degraded and sister chromatids migrate to opposing poles of the cell, thus enabling nuclear division (aka karyokinesis). Thereafter, actin redistributes to the contractile ring, which constricts creating the cleavage furrow. The cell cycle is completed by cytokinesis that splits the cytoplasm. Hence, the coordinated action of actin and microtubules is key to cell division. Errors in centrosome duplication, actin or microtubule polymerization, spindle assembly, or chromosome segregation lead to aneuploidy and/or mitotic catastrophe.

Formins are key regulators of actin dynamics. There are fifteen mammalian formins and they all possess two formin homolog (FH) domains (Breitsprecher and Goode, 2013). FH1 delivers profilin-bound actin monomers to the actin filament barbed ends accelerating elongation, whereas the FH2 dimerizes enabling the bundling of actin filaments (Kovar, 2006). Diaphanous (DIAPH) formins form a subgroup of 3 members that have three regulatory domains: a GTPase binding domain (GBD) at the N-terminus, a diaphanous autoregulatory domain (DAD), and a diaphanous inhibitory domain (DID). DAD binds to DID maintaining the DIAPH proteins in an inactive state. Activation occurs through the binding of Rho-GTPases, which releases the DAD/DID interaction. The best-characterized functions of DIAPH3 (also referred to as mDia2) are actin-related. In proliferating cells, DIAPH3 is required for the formation of the contractile ring and the cleavage furrow to enable cytokinesis (Chen et al., 2017, DeWard and Alberts, 2009, Watanabe et al., 2013). DIAPH3 is also involved in filopodia assembly, supporting cell migration (Stastna et al., 2012), and mesenchymal-amoeboid transition (Hager et al., 2012, Morley et al., 2015). In addition to the above-mentioned roles, evidence for DIAPH3 implication in actin/cytokinesis-independent functions has emerged. Early *in vitro* studies have suggested that DIAPH3 interacts with EB1 and APC, which localizes to plus-end, thereby stabilizing microtubules (Wen et al., 2004, Bartolini et al., 2008). Analysis of the full *Diaph3* knockout mice has revealed an important role of DIAPH3 in the biology of NSCs. The lack of DIAPH3 severely compromises chromosome segregation, leading to aneuploidy, mitotic catastrophe, and loss of these cells (Damiani et al., 2016). Yet, the precise function of DIAPH3 in nuclear division, especially in cytoskeletal rearrangements remains elusive. Furthermore, the impact of DIAPH3 deficiency on brain development and function were not assessed due to early embryonic lethality.

Here, we report that DIAPH3 localizes at the centrosome during mitosis and regulates the assembly of microtubules as well as the bipolar shape and orientation of the mitotic spindles. DIAPH3-deficiency disrupts the expression and/or stability of several proteins leading to multipolar spindles and disorganized cytoskeleton. One of the affected proteins is SPAG5 (also known as the mitotic spindle-associated protein 126 or Astrin), which localizes at the centrosome and kinetochore and has a documented function in cell division. SPAG5 displays a similar expression pattern as DIAPH3 in the developing cortex. Downregulation of SPAG5 phenocopies the DIAPH3 deficiency, whereas its overexpression rescues the DIAPH3 knockdown phenotype. We also used a conditional approach to delete *Diaph3* specifically in the cerebral cortex and report that this causes a marked depletion of cortical neurons, microcephaly, locomotor impairment, and social interaction defects.

## Results

### DIAPH3 localizes to centrosome and is required for assembly and function of mitotic spindle

We studied the subcellular distribution of DIAPH3 by immunofluorescence in U2OS cells. The protein co-localized with the centrosomal marker γ-tubulin during the whole mitosis (Fig. 1A-D). In telophase, it was also seen at the midzone (Fig. 1D), which is consistent with its documented role in the ingression of the cleavage furrow during cytokinesis (Watanabe et al., 2013, Damiani et al., 2016, DeWard and Alberts, 2009). We downregulated the expression of DIAPH3 by shRNA (Fig. 1E, F), and found that this disrupts cell division (82% of Diaph3-knockdown cells showed abnormal division versus 4% in control cells; Fig. 1G-J, and Supplementary information, Table S1). Notably, DIAPH3 knockdown compromised the integrity of the centrosome and bipolar shape of the spindle (40% of cells exhibited abnormal number of centrosomes); as well as the organization of microtubules, leading to mis-segregation of chromosomes (Supplementary information, Table S1). Knockdown of DIAPH3 also reduced cell survival to 59.1±0.73% of control cells (Supplementary information, Fig. S1A), likely by inducing apoptosis (percentage of aCas3+ cells: 1.1%±0.12% for scrambled sh-RNA versus 3.2%±0.16% for Diaph3 sh-RNA, Supplementary information, Fig. S1B). However, it did not affect cell proliferation as the ratio of Ki67+ cells was preserved (Supplementary information, Fig. S1C).

**Figure 1:**
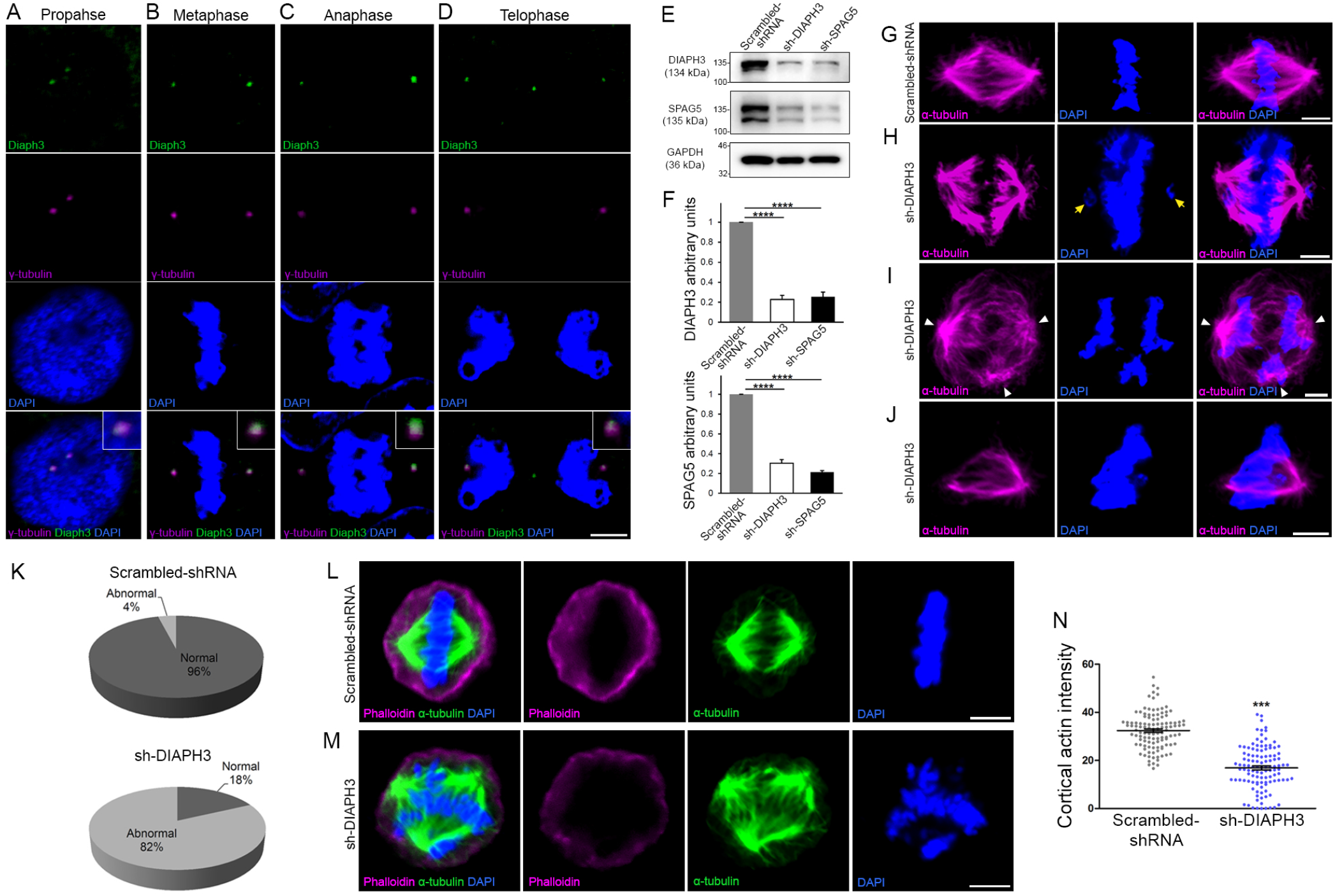
Loss of DIAPH3 disrupts the expression/stability of SPAG5 and causes mitotic defects. (**A-D**) Immunostaining of U2OS cells at prophase (**A**), metaphase (**B**), anaphase (**C**), and telophase (**D**), with anti-DIAPH3 (green) and anti-γ tubulin (magenta). The chromosomes were counterstained with DAPI (blue). DIAPH3 localizes at the centrosome at all mitotic stages. Insets are zooms in the centrosomal region showing that the DIAPH3 signal is pericentrosomal. Scale bar, 10 µm. (**E, F**) Western blot analysis of DIAPH3 and SPAG5 levels upon shRNA downregulation in U2OS cells. shRNA against DIAPH3 (sh-DIAPH3) significantly reduced levels of DIAPH3 and SPAG5 proteins (*P*= 0.000044, and 0.000046 respectively). Reciprocally, shRNA against SPAG5 (sh-SPAG5) reduced SPAG5 and DIAPH3 protein levels (*P*= 0.000001, and 0.000097 respectively). Scrambled-shRNA were used as control for the transfection and GAPDH as loading control. *n*=3 independent experiments, Student’s *t*-test, error bars represent s.e.m. (**G-J**) Representative images of mitotic cells transfected with scrambled-shRNA (**G**) and shRNA against DIAPH3 (sh-DIAPH3) (**H-J**), immunostained with anti-α tubulin (magenta). Chromosomes were counterstained with DAPI (blue). Yellow arrows in (**H**) point to lagging chromosomes. Arrowheads in (**I**) depict poles of a mitotic spindle. (**J**) Representative image of a mitotic cell with disorganized/shrunk microtubules, single centrosome/pole, and asymmetric metaphase plate. Scale bar, 5 µm. (**K**) quantification of mitotic errors. 82% of DIAPH3 Knockdown cells exhibit mitotic abnormalities (4% in control cells). *n*=118 and 115 cells from 5 individual experiments of scrambled-shRNA and sh-DIAPH3 respectively. (**L-N**) Diminished cortical actin in DIAPH3 knockdown cells. (**L, M**) Illustration of mitotic cells transfected with scrambled-shRNA (**L**) or sh-DIAPH3 (**M**), and immunostained with phalloidin (magenta) and α-tubulin (green). Chromosomes were counterstained with DAPI; Scale bar, 5 µm. (**N**) Quantification of cortical actin (fluorescence intensity) showing a reduction of 48% of cortical actin in DIAPH3 knockdown cells. *n*=113 cells per condition from 3 distinct experiments. Student’s *t*-test, *P*=1.07×10^−32^. Error bars represent s.e.m.

### Downregulation of DIAPH3 alters SPAG5 expression

Positioning of the spindle during mitosis is governed by multiple mechanisms. Key among those is the “cortical pulling”, which refers to the capacity of specific sites on the cell cortex to capture and exert forces on astral microtubules to orient the mitotic spindle. Cortical pulling is steered by cytocortical proteins that involve nuclear mitotic apparatus (NUMA), G-protein signaling modulator 2 (GPSM2, also known as LGN or partner of inscrutable (PINS)); polarity proteins PAR3, NUMB; and the adapter protein Inscuteable (INSC). The assembly of NUMA-GPSM2 beneath the cortical actin, recruits the dynein/dynactin motor protein complex to haul astral microtubules (Morin and Bellaiche, 2011). Downregulation of DIAPH3 dramatically reduced the amount of cortical actin (Fig. 1L-N). We used Western blotting on telencephalic lysates from *Diaph3* KO mice (Damiani et al., 2016) to assess the level of core cortical proteins NUMA, GPSM2, PAR3, NUMB, INSC, dynein, dynactin; microtubules plus-end associated proteins sperm associated antigen (SPAG 5), Kinastrin (KNSTRN, aka small kinetochore-associated protein/SKAP); and the cytoplasmic linker associated protein (CLASP)1. We also used CENPA, a protein downregulated in DIAPH3 depleted cells (Liu and Mao, 2016) as positive control for the Western blotting and normalized the results to GAPDH levels. We did not detect significant changes in the level of endogenous NUMA, dynein, dynactin, INSC, or CLASP1 between KO and control mice. In contrast, polarity proteins GPSM2, NUMB, PAR3, and microtubule associated proteins SPAG5 and KNSTRN were downregulated (Fig. 2A, B and Supplementary information, Table S2). The most prominent reduction was observed in SPAG5. We confirmed this by real time RT-PCR, which showed a 75% reduction of the transcript (Fig. 2C), and by RNAscope situ hybridization (Fig. 2D). The expression pattern of *Spag5* resembled that of *Diaph3* with a strong expression in cortical progenitor cells (Fig. 2D, see also http://www.eurexpress.org/ee/databases/assay.jsp?assayID=euxassay_004999&image=01, and (Damiani et al., 2016). To explore further the relationship between DIAPH3 and SPAG5, we knocked-down the former in U2OS cells, and used immunofluorescence to scrutinize the expression of the latter. In control cells (transfected with scrambled-shRNA), SPAG5 localized to the spindle microtubules and kinetochore during metaphase and progressively concentrated in the pericentrosomal region and spindle poles as the cell proceeds to anaphase (Fig. 2E, F). Importantly, SPAG5 expression declined sharply in DIAPH3-depleted cells (Fig. 1E, F and Fig. 2G-J), and its distribution was disrupted (Fig. 2G-J, and Supplementary information, Table S1). In these cells, the bipolar organization of the spindle was altered, and astral microtubules were shrunk. The chromosomes failed to congress at the metaphase plate and did not migrate to the cell poles at anaphase (Fig. 2G-J).

**Figure 2:**
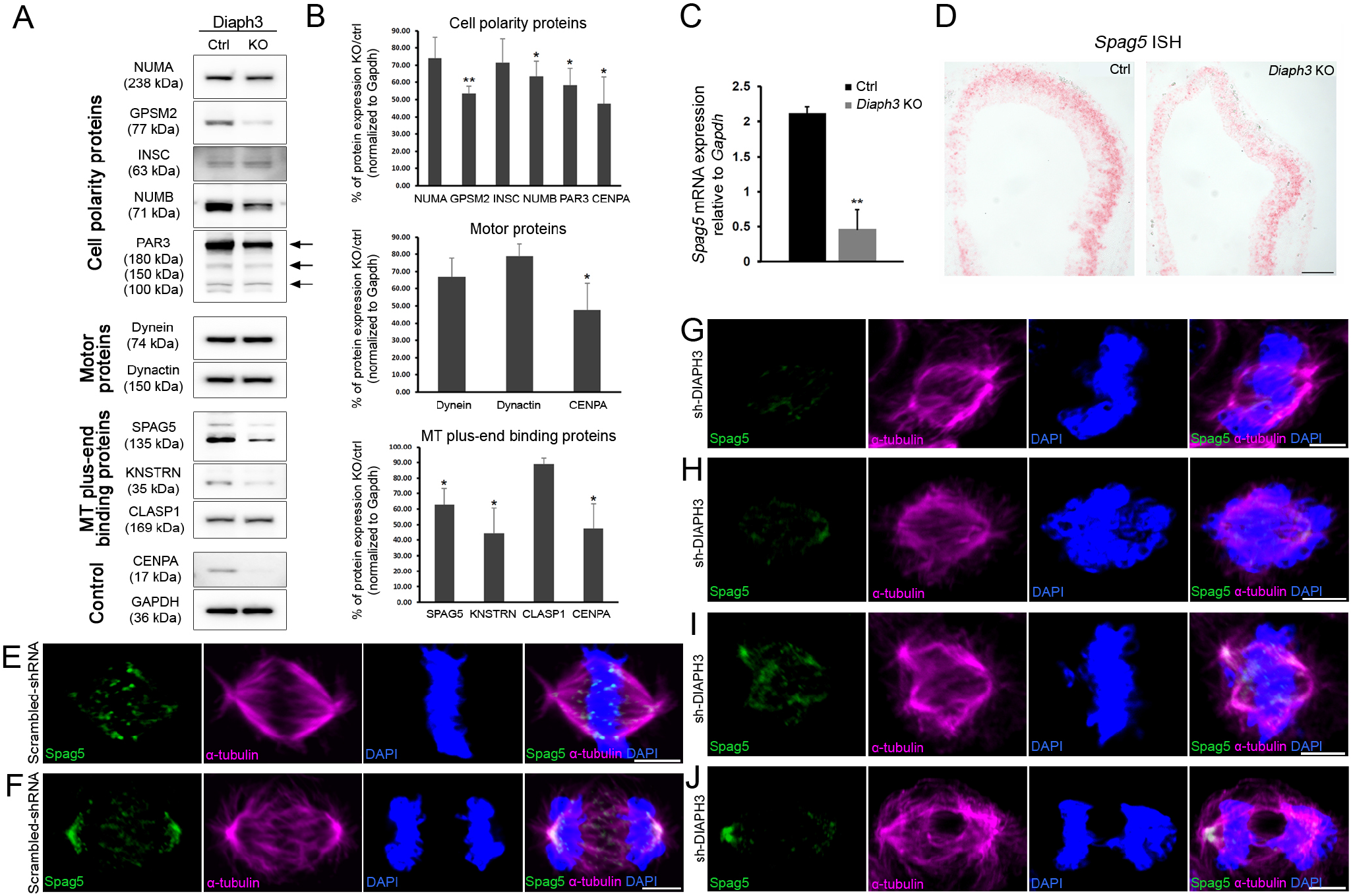
Diaph3-deficiency impairs the expression and stability of mitotic spindle polarity proteins. (**A, B**) Assessment of NUMA, GPSM2, INSC, NUMB, PAR3, Dynein, Dynactin, SPAG5, KNSTRN, and CLASP1 levels in telencephalon extracts of *Diaph3* KO mice by western blotting. CENPA was used as positive control (Liu and Mao, 2016) and GAPDH as loading control for quantification (*n*=3 embryos for each genotype. *: *P*<0.05, **: *P*<0.01, Student’s t-test, error bars represent s.e.m. Fold change and *P-*value are listed in Supplementary information, Table S2. (**C**) Quantification of the *Spag5* mRNA by real time RT-PCR. There is a reduction of 78% in *Diaph3* KO relative to control mice (*P*=0.0046). (**D**) Coronal sections of e11.5 Forebrain from control (left) and *Diaph3* KO (right, hybridized with a *Spag5* fast red-labelled RNAscope probe. The expression of *Spag5* mRNA is downregulated in *Diaph3* KO; scale bar, 150 µm. (**E-J**) Representative images of mitotic cells transfected with scrambled shRNA (**E**, **F**), or DIAPH3 shRNA (sh-DIAPH3, **G**-**J**), and immunostained with anti-SPAG5 (green), and anti-α tubulin (magenta) antibodies. Chromosomes were counterstained with DAPI (blue). SPAG5 localized at the centrosome and kinetochore in control cells (**E**, **F**), and both its expression level and distribution were altered in DIAPH3 knockdown cells (**G-J**). Scale bar, 5 μm.

### Downregulation of SPAG5 phenocopies the DIAPH3 phenotype

SPAG5 is an essential component of the mitotic spindle. It is required for chromosome alignment, sister chromatid segregation, and progression to anaphase (Mack and Compton, 2001, Gruber et al., 2002, Thein et al., 2007). SPAG5 promotes microtubule-kinetochore attachments, and regulates the localization of several centrosomal-proteins (e.g. CDK2, CDK5RAP2, CEP152, WDR62, and CEP63) (Kodani et al., 2015). Silencing SPAG5 in U2OS cells triggered multipolar spindles and chromosome mis-segregation, a phenotype reminiscent of the DIAPH3 knockdown (Fig. 3A-C) (Thein et al., 2007). It also compromised the expression and/or stability of DIAPH3 (please compare Fig. 3A-C with Fig. 1B, C). Remarkably, overexpression of SPAG5 significantly rescued the DIAPH3 knockdown phenotype and partially restored cell viability (survival: 87.1±1.71% in SPAG5 rescue versus 59.1±0.73% in DIAPH3 knockdown; aCas3+ cells: 1.1±0.15% in SPAG5 rescue versus 3.2±0.16% in DIAPH3 knockdown; Fig. S1). These results along with the expression of SPAG5 and its downregulation in the *Diaph3* KO mice prompted us to investigate its function *in vivo*. We electroporated e13.5 embryos with scrambled-shRNA or sh-SPAG5 cDNA constructs and analyzed cell division of aNPC at e15.5. We observed multipolar spindles and a bias towards asymmetric/neurogenic divisions in brains electroporated with sh-SPAG5 (Fig. 3D-G). 49% (67/137) of SPAG5 knockdown cells divide asymmetrically versus 31% (45/147) in control cells (Fig. 3H, I).

**Figure 3:**
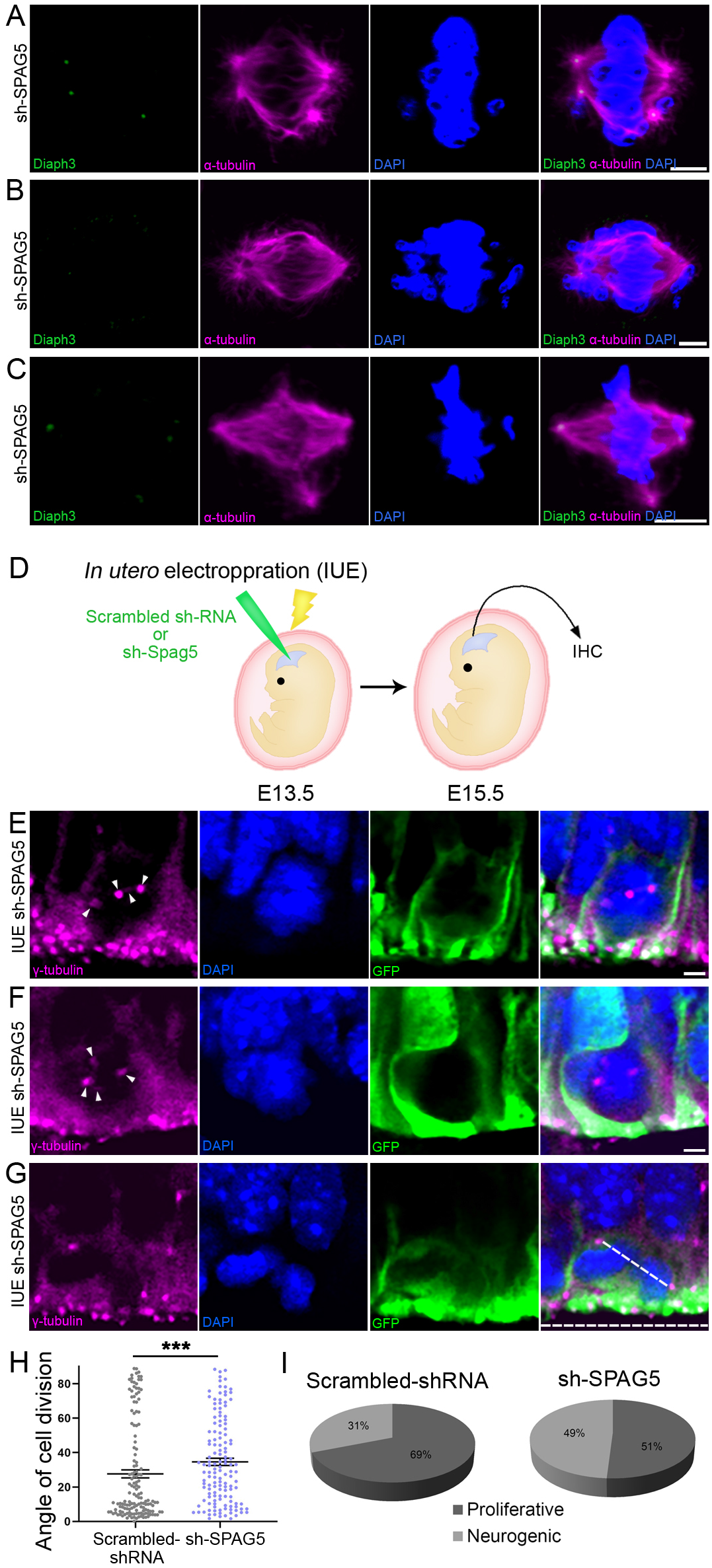
Downregulation of SPAG5 phenocopies DIAPH3 deficiency. (**A-C**) Representative images of mitotic cells transfected with SPAG5 shRNA (sh-SPAG5) and immunostained with anti-DIAPH3 (green) and anti-α tubulin (magenta) antibodies. Chromosomes were counterstained with DAPI (blue). SPAG5 deficiency downregulated DIAPH3 and caused mitotic errors. Note the mis-localization of DIAPH3 and spindle abnormalities in sh-SPAG5 transfected cells. Scale bar, 5 µm. (**D-H**) schematic representation of *in utero* electroporation of sh-SPAG5 in cortical progenitors at e13.5 and immunohistochemistry (IHC) analysis at e15.5. (**E, G**) Ventricular zone of telencephalic sections of e15.5 embryos electroporated *in utero* with SPAG5 shRNA and GFP at e13.5, and immunostained with anti-γ tubulin (magenta) at E15.5. White arrowheads in (**E** and **F**) depict numerical abnormalities of the centrosome. (**G**) Illustration of a neural progenitor undergoing neurogenic/asymmetric division. The horizontal and oblique dashed lines delineate the ventricular surface and mitotic spindle orientation, respectively. Scale bar = 2 µm. (**H, I**) Quantification of cell division modalities in electroporated aNPC. There is a bias towards neurogenic divisions (large angles) in sh-SPAG5 electroporated aNPC when compared with scrambled-shRNA, *P*=0.0009. Student’s *t*-test, error bars represent s.e.m. (**H**). 49% (67/137 cells) of sh-SPAG5 progenitors undergo neurogenic/asymmetric division compared to 31% (45/147 cells) in scrambled-shRNA (**I**).

### Cortex-specific inactivation of *Diaph3* disrupts neurogenesis

The lack of DIAPH3 compromises nuclear division in neural stem cells (NSC), and induce apoptosis (Damiani et al., 2016). To evaluate the fraction of dying cells, we performed aCas3 immunostaining at E11.5. Around 18% of NSC were aCas3 positive in *Diaph3 KO* mice (18±0.83% of cells in KO versus 2.4±0.65% in control; Supplementary information, Fig. S2). To assess the effect of DIAPH3 loss on cortical histogenesis, we produced conditional knockout mice (cKO) in which, we specifically inactivated *Diaph3* in the cerebral cortex by crossing a floxed allele with mice expressing the recombinase Cre under the control of *Emx1* promoter (Supplementary information, Fig. S3). We quantified the number of aNPC (Pax6), BP (Tbr2), and neurons (Tbr1) at E13.5, and found that the three populations were affected (Fig. 4A-I). Their respective numbers decreased by 36% (number of Pax6^+^ cells/0.1mm^2^= 502±48 in cKO, versus 786±35 in control, Fig. 4A-C), 31% (number of Tbr2^+^ cells/0.1mm^2^= 234±4.8 in cKO, versus 339±15 in control Fig.4D-F), and 23% (number of Tbr1^+^ cells/0.1mm^2^= 404±12 in cKO, versus 524±18 in controls, Fig. 4G-I). These results show that the loss of DIAPH3 has an effect on the three cortical cell types with aNPC being the most affected population. BP and neurons are produced by neurogenic division, and settle at more basal positions than their precursors. This type of division, supposedly involves asymmetric inheritance of fate determinants, and has been correlated with a mitotic spindle that is perpendicular (cleavage plane horizontal) to the ventricular surface (Chenn and McConnell, 1995, Shitamukai et al., 2011). In this configuration, the cellular components and/or molecules regulating fate decision (e.g. the apical process, mother *vs* daughter centriole, cilia remnants, or polarity proteins) are unequally inherited by the daughter cells, even though this is still debated (Noctor et al., 2008). Given the deregulation of polarity proteins NUMB, and PAR upon DIAPH3 downregulation, we analyzed the mitotic spindle orientation in dividing aNPC *in vivo*. We used γ-tubulin (centrosome) and DAPI (chromosomes) to envision the orientation of the spindle at anaphase, and assess the mode of cell division (Fig. 4J). We considered dividing cells in both VZ and SVZ. In control embryos, 87 % (77/89) of divisions were symmetric (proliferative) and 13% (12/89) were asymmetric (neurogenic). In the *cKO*, the fraction of proliferative divisions dropped to 46% (37/81), whereas that of neurogenic divisions increased to 54% (44/81) (Fig. 4K, L). We also observed signs of multipolar spindles in cKO tissue (Fig. 4M, N), a phenotype that is reminiscent of DIAPH3 and SPAG5 knockdown (Fig. 1 and Fig. 3).

**Figure 4:**
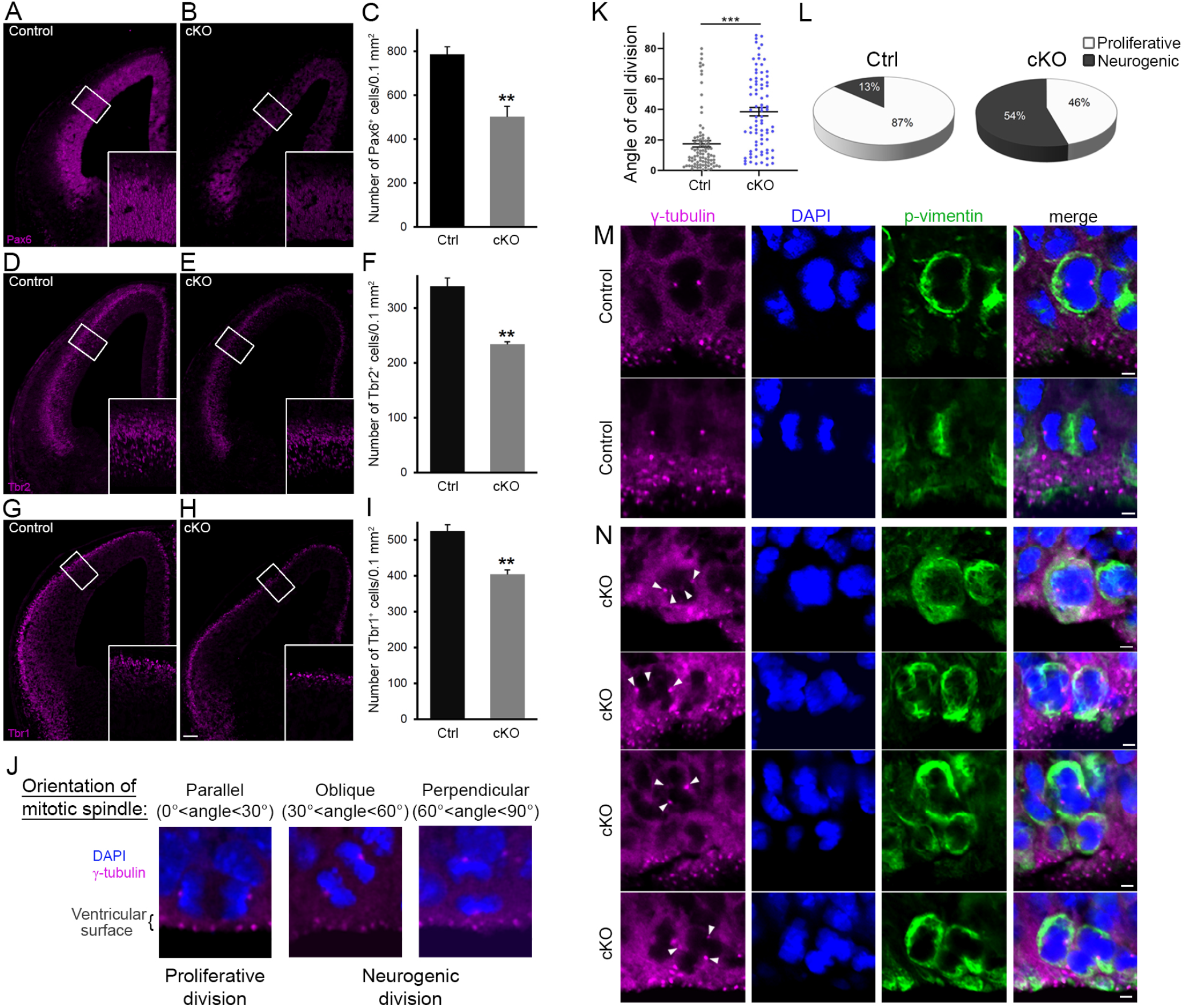
Cortex-specific inactivation of *Diaph3* disrupts cortical neurogenesis. (**A, B, D, E, G,** and **H**) Forebrain coronal sections from e13.5 stained with Pax6 (aNPC, **A, B**), Tbr2 (basal progenitors, **D, E**) and Tbr1 (neurons, **G, H**) antibodies. Insets are enlargements of the boxed areas. Quantifications shown in (**C, F,** and **I**), emphasize a reduction in the number of apical radial glia (aNPC, Pax6^+^, *P*=0.00853), basal progenitors (Tbr2^+^, *P*=0.00287), and neurons (Tbr1^+^, *P*=0.00535) respectively. Cells were counted in 0.1 mm^2^ cortical areas. *n*=3 embryos per genotype. Student’s *t*-test, error bars represent s.e.m. Scale bar, 100 µm. (**J**) e12.5 cortical sections stained with anti-γ tubulin antibodies (magenta) and DAPI (blue) to label centrosomes and chromosomes respectively and “foresee” the mitotic spindle. The orientation of the mitotic spindle with respect to the ventricular surface is categorized into 3 types: parallel (left), oblique (middle), or perpendicular (right). When the mitotic spindle is parallel to the ventricular surface (0°<angle<30°), the cell division is proliferative. When it is oblique (30°<angle<60°) or perpendicular (60°<angle<90°) to the ventricular surface, the division is neurogenic. (**K, L**) Assessment of cell division modality at e12.5. Compared with control mice, there is a significant shift toward larger angles (*P*=5.9×10^−9^, *n*=89 mitoses from 3 control embryos, 81 mitoses from 3 cKO embryos. Student’s *t*-test, error bars represent s.e.m) (**K**); and an increase in the ratio of neurogenic division at the expense of proliferation in cKO (**L**). (**M**) Dividing aNPC in control telencephalic VZ stained with anti-γ tubulin antibodies (magenta), anti-phospho vimentin antibodies (green) and DAPI (blue) to label centrosomes, dividing cells and chromosomes respectively. (**N**) Illustrations of supernumerary centrosomes (arrowheads) in cKO. Scale bar, 2 µm.

### *Diaph3* cKO mice exhibit cortical hypoplasia and impaired behavior

The cerebral cortex of adult *Diaph3* cKO mice was markedly thin (Fig. 5A, B). Immunostaining with markers of different cortical layers (Cux1: layer II-III, Foxp2: layer VI) revealed a diminished number of cells along with a reduced thickness of cortex (Fig. 5C-F). We then assessed the consequences of cortical hypoplasia in terms of behavior. cKO mice displayed lower spontaneous locomotor activity in open-field test and elevated plus maze (Fig. 5G, H), but no difference was detected when we analyzed the time spent in the periphery of the open-field, or in the closed arms of the elevated plus maze (Supplementary information, Fig. S4A, B). These results suggest that DIAPH3 deficiency impairs locomotor activity, but has little effect, if any, on anxiety and/or attention. In addition, the cKO mice exhibited a defective social behavior in the “three-chamber” test. They spent more time in empty chamber than with a stranger mouse (stranger 1), emphasizing a lower sociability of *cKO* mice, as compared to control littermates (Fig. 5I). When another stranger mouse was introduced, the *cKO* mice spent less time with the new stranger, indicating impairment in social novelty behavior (Fig. 5J). cKO mice did not show any olfactory defect (Supplementary information, Fig. S4C). They performed as well as controls in the Morris water maze (Supplementary information, Fig. S4D), and Y-maze tests (Supplementary information, Fig. S4E), implying that long-term and short-term memories were preserved. Finally, they did not display any repetitive behavior as assayed by self-grooming (Supplementary information, Fig. S4F), or marble burying (Supplementary information, Fig. S4G). These results show that the cKO mice have specific deficits in motor activity and social behavior which is in line with previous studies that have associated DIAPH3 mutations with autism spectrum disorder in humans (Vorstman et al., 2011, Xie et al., 2016).

**Figure 5:**
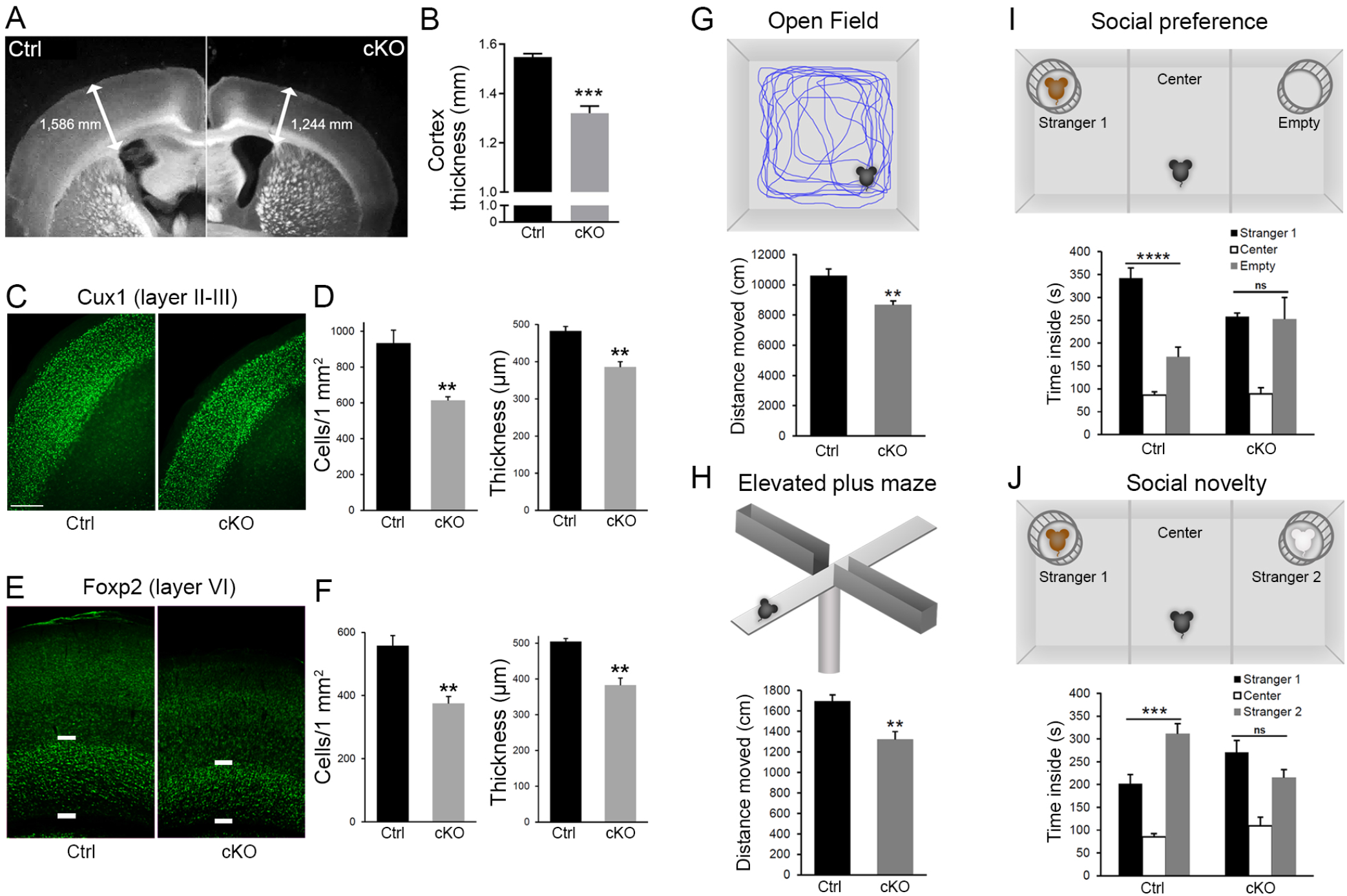
*Diaph3* cKO mice display cortical hypoplasia and behavioral defects. (**A**) Dark field micrograph of coronal sections of the forebrain from control (left) and cKO (right) mice, depicting a marked reduction of the cortex thickness in cKO; quantified in (**B**) (1.3±0.028 mm in cKO versus 1.5±0.014 mm in control, *P*=0.00038). (**C**) Coronal sections stained with the upper layer marker Cux1. (**D**) Quantification of the number of Cux1^+^ cells (left) and thickness (right) in layers II-III (Cux1^+^ cells: 614±20 cells in cKO versus 933±73 in control, *P*=0.0056; thickness: 385±15 μm in cKO versus 482±13 μm in control, *P*=0.0025). (**E**) Coronal sections stained with deep layer marker Foxp2. (**F**) Quantification of the number of Foxp2^+^ cells (left) and thickness (right) in layer VI (Foxp2^+^ cells: 375±22 cells in cKO, versus 558±32 in control, *P*=0.0033; thickness: 383±20 μm in cKO versus 505±8 μm in control, *P*=0.0012). Cux1^+^ and Foxp2^+^ cells were counted in 1 mm^2^ cortical area. *n*=4 embryos for each genotype, Student’s *t*-test, error bars represent s.e.m. Scale bar, 200 µm. (**G**) Distance moved by cKO and control mice in the open field. Student’s *t* test, *P*=0.0015. (**H**) Distance travelled by mice in elevated plus maze. Student’s *t* test, *P*=0.0043. (**I**) Social behavior in the “3 chambers” test. One-way ANOVA test, *P*=8.4×10^−7^ for control, *P*=0.99 for cKO (stranger versus empty chamber). (**J**) Social novelty behavior in the “3 chambers” test. One-way ANOVA test, *P*=0.00050 for control, *P*=0.16 for cKO (stranger 1 versus stranger 2 chamber). Compared with control, cKO mice have a reduced locomotor activity and defective social interactions. *n*=10 per genotype. Error bars represent s.e.m.

## Discussion

DIAPH3 is an effector of Rho GTPases that has been classically associated with actin dynamics *in vitro* (Watanabe et al., 2010, Watanabe et al., 2013). Functional analysis of knockout mice showed that it regulates cytokinesis in erythroid cells by promoting the accumulation of actin in the cleavage furrow at telophase (Watanabe et al., 2013). In addition to this well-documented role in actin cytoskeleton and cytokinesis, we have reported that DIAPH3 plays a role during karyokinesis, and that its loss causes chromosome mis-segregation, aneuploidy, and cell death (Damiani et al., 2016). Here we provide evidence that DIAPH3 localizes to the centrosome during mitosis. Its depletion compromises the integrity of centrosome, spindle and astral microtubules, as well as the expression or stability of several proteins involved in spindle-cell cortex and spindle-kinetochore interactions (Fig. S5). These results are consistent with previous *in vitro* data suggesting that DIAPH3 may serve as scaffold protein that binds to, and stabilizes microtubules (Palazzo et al., 2001, Bartolini et al., 2008, Wen et al., 2004). By regulating the centrosome integrity, microtubule stability and spindle orientation, DIAPH3 sits at the heart of cell division accuracy and fate determination (Fig. S5). All these processes are requisites for the production and maintenance of neural progenitors and neurons. The centrosome acts as a seed and anchoring point for the minus end of microtubules as they grow toward the equatorial region (spindle microtubules) or the cell cortex (astral microtubules). Consistent with this, we found that depletion of DIAPH3 disrupts the number of centrosomes, which results in disorganized spindle and astral microtubules, abnormal alignment and segregation of chromosomes. DIAPH3 depletion affects the expression level and/or distribution of several proteins. Among those, SPAG5 which localizes at centrosome and kinetochore, is essential to the recruitment of other centrosomal proteins such as CDK5RAP2 (MCPH3) (Kodani et al., 2015). SPAG5 controls sister chromatid cohesion by regulating separase activity (Thein et al., 2007). Knockdown of SPAG5 induces similar mitotic errors to those triggered by knockdown of DIAPH3 ((Kodani et al., 2015, Thein et al., 2007) and this work), and its overexpression rescues the DIAPH3 knockdown’s phenotype. DIAPH3 depletion also affects the expression level or the stability of GPSM2, NUMB and PAR3, all of which localize at the region of the cell cortex where astral microtubules are anchored.

Microcephaly is a reduction of more than 3 standard deviations of the head circumference compared to matching gender, age and ethnicity controls (Kaindl et al., 2010, Jayaraman et al., 2018). Primary microcephaly is an inherited neurodevelopmental defect that appears mostly during pregnancy or at the first postnatal year. Eighteen “microcephaly” genes named MCPH 1-18 (for microcephaly primary hereditary 1-18) have been identified (Jayaraman et al., 2018). All these genes are expressed in the germinal zones wherein neural progenitors reside, proliferate, and differentiate into neurons during the second trimester of pregnancy (Verloes et al., 1993, Gilmore and Walsh, 2013). A key determinant of the brain size is the production and maintenance of the right numbers of neural progenitors and neurons. To attain the initial pool of progenitors and right number of neurons, neural stem cells must reconcile two imperatives: the high speed and fidelity of cell division. This relies on the interplay between cell cycle regulators, mitotic spindle assembly, mitotic checkpoints, and cytoskeleton. Hence, division of neural stem cells and the balance between proliferation and differentiation of apical progenitors is a critical factor of the brain size (Gilmore and Walsh, 2013). Whether an apical progenitor undergoes a proliferative or neurogenic division depends on inheritance of polarity components, which has been correlated with the cleavage plane and orientation of the mitotic spindle. If the cleavage plane is parallel to the apical-basal polarity axis (spindle parallel to the ventricular surface), the two daughter cells are likely to inherit equally the cell components, giving rise to two identical progenitors. In contrast, when the cleavage plane is perpendicular or oblique to the apical-basal axis, the two daughter cells asymmetrically inherit apical and basal components, which gives rise to one apical progenitor and one neuron, or one apical and one basal progenitor. Hence, defects in the balance of proliferative versus neurogenic division or in the timing of the neurogenic switch affect cell fate and the production of neurons.

Loss of function of DIAPH3 disrupts the modality of cell division and fate determination of apical progenitors, enhancing the number of neurogenic/asymmetric division at the expense of proliferative/symmetric division, and prematurely exhausting the pool of progenitors. Even though the cytocortical proteins help to orientate the mitotic spindle, the loss of GPSM2 does not lead to microcephaly despite randomized cleavage plane in dividing neural progenitors (Konno et al., 2008, Blumer et al., 2008). Therefore, the role of DIAPH3 in fidelity of nuclear division and the cell loss observed in *Diaph3* KO mice may be more instrumental to emergence of microcephaly in cKO mice than its role in oriented cell division. It will be interesting to test whether the blockade of apoptosis induce neoplastic transformation due to accumulation of aneuploid cells. Finally, DIAPH3 is expressed in several epithelia and may govern cell proliferation, therefore influencing the size of different organs and/or organisms. In support of this, patient with microcephalic osteodysplastic primordial dwarfism type 1 (MOPD 1) syndrome display a 50% reduction in DIAPH3 expression (Edery et al., 2011). Furthermore, *Diaph3* has been associated with body size in equines, as copy number variants were reported in donkeys, ponies, and horses with short stature (Metzger et al., 2018).

In humans, two clinical studies point to *DIAPH3* as an autism susceptibility gene (Vorstman et al., 2011, Xie et al., 2016). The first study identified a double hit mutation with one amino acid substitution (Pro614Thr) in one allele, and a deletion in the second allele. The second is a *de novo* point mutation in the 18th exon (c.2156T>C, pI719T). Several chromosomal deletions spanning the *DIAPH3* locus have also been associated with language impairment, autism and intellectual disability (https://decipher.sanger.ac.uk/, and (Nathalie Sans, 2016)). Importantly, among autistic patients, a significant fraction exhibit microcephaly (Fombonne et al., 1999). In this work, we report that the conditional deletion of *Diaph3* in the mouse cerebral cortex dramatically affects neurogenesis, leads to microcephaly and autistic-like behavior, with typical altered motor activity and social interactions. Our finding that the deletion of *Diaph3* affects the accuracy and modality of neural progenitor division ultimately regulating the final number of cortical neurons, may provide a molecular and cellular basis to the brain malformations and neurological disorders that have been associated with DIAPH3 dysfunction in humans.

Combined together, our results suggest a novel role of DIAPH3 as a centrosomal protein. Its deficiency induces supernumerary centrosomes, and disrupts spindle and astral microtubule. These defects cause inaccuracies in nuclear division and fate decision of neural progenitors, leading to the loss of cortical progenitors, microcephaly, and autistic-like behavior.

## Methods

### Mutant mice

All animal procedures were carried out in accordance with European guidelines and approved by the animal ethics committee of the Université Catholique de Louvain. Mouse lines used in this study were: *Emx1^tm1(cre)Krj^/J*; Jackson Lab) (Gorski et al., 2002, Damiani et al., 2016), *Diaph3 KO*, *Diaph3*^*Emx1-Cre*^*cKO* (*Diaph3*^*f/f*^; *Emx1-Cre*), and *Diaph3*^*f/f*^ (control for the cKO).

### Immunostaining and antibodies

For immunohistochemistry, embryos were fixed in 4% paraformaldehyde (PFA), cryoprotected by gradients of sucrose solution. Cryosections were blocked in PBST (0.1% Triton X-100 in PBS) supplemented with 5% normal goat serum and 1% BSA for 30 min. Slides were incubated in primary antibodies diluted in blocking buffer at 4°C overnight. Slides were washed and incubated with secondary antibodies (Alexa, Thermofisher) diluted in the same blocking buffer. Cultured cells were fixed with chilled methanol for 5 min and then followed by washing in PBST. Cells were incubated in blocking buffer for 30 min, primary antibodies for 1.5 h and secondary antibodies for 1.5 h at room temperature. Primary antibodies were as follow: rabbit anti-Cux1 (Santa Cruz, SC-13024, 1:200), rabbit anti-Foxp2 (Abcam, ab16046, 1:500), rabbit anti-Tbr1 (Abcam, ab31940, 1:500), rabbit anti-Tbr2 (Abcam, ab23345, 1:500), rabbit anti-Pax6 (Covance, PRB-278P, 1:500), rabbit anti-γ-tubulin (Abcam, ab11317, 1:500), Diaph3 (1:500) (Tominaga et al., 2000), rabbit anti-Spag5 (Proteintech, 14726-1-AP, 1:200), mouse anti-α-tubulin (Sigma, T6199, 1:500), rabbit anti-aCas3 (Cell signaling, 9603, 1:100), and mouse anti-Ki67 (BD phamingen, 556003,1:500). Nuclei/chromosomes were counterstained with DAPI. Images were acquired with an Olympus FV1000 confocal microscope. Mitotic cells were imaged with Z-stack that focused on the levels of centrosomes.

### Western blotting

Tissues were homogenized in lysis buffer containing 50 mM Tris HCl pH 7.5, 150 mM NaCl, 1% NP40 and protease inhibitors (Roche). Cell lysates were incubated on ice for 30 min prior to centrifugation at 2000g for 10 min at 4°C. Protein quantification was performed with BCA Protein Assay kit (Pierce). Supernatant were mixed with SDS-loading buffer and heated at 85°C for 10 min. Equal amount of proteins were loaded on 4-12% gel (Invitrogen) and transferred to PVDF membrane (Merck Millipore). Membranes were blocked with StartingBlock buffer (ThermoScientific) and incubated overnight at 4°C with rabbit anti-DIAPH3 (1:5000) (Tominaga et al., 2000), rabbit anti-SPAG5 (Sigma, HPA022008, 1:750;), chicken anti-GAPDH (Millipore, AB2302, 1:2000), mouse anti-NUMA (Becton Dickinson, 610561, 1:500), rabbit anti-PAR3 (Millipore, 07-330, 1 :500), rabbit anti-GPSM2 (1 :200) (Ezan et al., 2013), rabbit anti-CENPA (Cell Signaling, 2048, 1 :500), mouse anti-dynein (Santa Cruz, sc-13524, 1 :500), rabbit anti-KNSTRN (Sigma, HPA042027, 1:1000), rabbit anti-INSC (Abcam, ab102953, 1:1000), goat anti-NUMB (Abcam, ab4147, 1:100), mouse anti-dynactin (BD Biosciences, 610474, 1:500), rabbit anti-CLASP1 (Abcam, ab108620, 1:5000). Proteins were detected with SuperSignal West Pico PLUS solution (ThermoScientific) or SuperSignal West Femto Maximum Sensitivity Substrate kit (ThermoScientific). Band intensity was imaged by fusion pulse and quantified with ImageJ Gel analyzer tool. Values were normalized with GAPDH. The amount of protein in control was set to one.

### qPCR

Total mRNA was isolated from control or *Diaph3* KO E11.5 telecephalon using the RNeasy mini Kit (Qiagen) according to the supplier’s instructions. Reverse transcription was performed with an RT cDNA Synthesis Kit (Promega). Real-time PCR was performed with SYBR green SuperMix using an iCycler real-time PCR detection system (Bio-Rad).

### Cell culture and knockdown experiments

U2OS cell were cultured in DMEM supplemented with penicillin-streptomycin and 10% FBS (Invitrogen). Cells were seeded in 12-well plates (day 0) and were transfected with Lipofectamine LTX (Invitrogen) (day 1) using scrambled shRNA (Origene, TR30012) or pool of sh-Diaph3 (Origene, TR304992) with sequences: ATTTATGCGTTGTGGATTGAAAGAGATAT, AATCAGCATGAGAAGATTGAATTGGTTAA, CACGGCTCAGTGCTATTCTCTTTAAGCTT, AAGAGCAGGTGAACAACATCAAACCTGAC, or pool of sh-Spag5 (Origene, TR309161) with sequences: CCTCAAGGACACTGTAGAGAACCTAACGG, GGTAGGATTCTTGGCTCTGATACAGAGTC, CTCCAAGGAAAGCCTGAGCAGTAGAACTG, AGATGAAGAGCCAGAATCAACTCCTGTGC. Cells were harvested 72 h after transfection for western blotting. For immunostaining, cells were serum-deprivated at day 3 (48 h after transfection) and seeded onto glass coverslip pre-coated with gelatin at day 4 (72 h after transfection). Cell cycle was synchronized by serum add back, and cells were fixed after 16 h. For cell survival assay, cells were seeded at the same density (50% confluency, 2 × 10^5^ cells per well) onto glass coverslip pre-coated with gelatin in 12-well plates at day 0. Cell number was count at day 1 (just before transfection) and day 4 (72 h after transfection). Cells were then immunostained with apoptotic marker (aCas3), proliferation marker (Ki67) and DAPI. Cell density was imaged and counted by Zen lite with Zeiss AXIO light microscope.

### Behavioral tests

All the behavioral tests were performed using adult cKO *(Diaph3^Emx1-Cre^ cKO (Diaph3^f/f^*; *Emx1-Cre*), and control littermates (*Diaph3^f/f^*) males.

The “open-field” test was performed to assess locomotor activity and anxiety. Mice were allowed to move freely in a square arena (60 × 60 cm) and video tracked (Ethovision 6.1, Noldus; Wageningen, The Netherlands) for 20 min (Mallon et al., 2008, Mignion et al., 2013). The total distance covered by test mice (locomotor activity) and the time spent in the periphery (anxiety) were measured.

The “Elevated plus maze” test” was performed to assess locomotor activity and anxiety. Mice were placed in a setup consisting of two opposing open arm (exposed place) and two opposing closed arm (safer place). Total distance travelled and time spent in closed arm were recorded by a video tracking system (Ethovision 6.1, Noldus; Wageningen, The Netherlands) for 5 min.

The “Three-chamber” test was performed to assess sociability. Test box was divided into 3 equal compartments with 20 cm each and dividing walls had retractable doorways allowing access into each chamber. Mice were habituated in the middle chamber for 5 min with the doorways closed. To evaluate social interactions, mice were enclosed in the center compartment and an unfamiliar mouse (called stranger 1) was restricted in a wire cage placed in one side compartment while the other side compartment contains an empty wire cage. Mice were video tracked (Ethovision 6.1, Noldus; Wageningen, The Netherlands) for 10 min. The time spent in each chamber was measured. To evaluate the preference for social novelty, a second unfamiliar mouse (called stranger 2) is introduced into the empty wire cage in the other side compartment of the sociability test box. Mice were video tracked for 10 min and the time spent in each chamber was measured (Moy et al., 2007).

The “food localization” test was performed to evaluate the olfactory function. Test mice were fasted for 16 h with water supply ad libitum. They were then transferred into a clean cage with 3 cm thick of wood-chip bedding and lightly tamped down to make a flat surface. 1.0 g of food pellet was placed at the same location under the bedding and mice were introduced into the cage at a constant position. The time from which the mouse was placed into the cage until it retrieved the food pellet with its front paws was counted up to a maximum of 300 s.

The “Morris water maze” test was performed to assess long-term memory. Water maze was made of a round pool with a diameter of 113 cm virtually divided into four quadrants (North, South, West and East) and filled with water (26°C). Several visual cues were placed around the pool. The platform was placed at the center of the North-East quadrant of the pool and maintained in this position throughout the four days. Mice were video-tracked (Ethovision 6.1, Noldus; Wageningen, The Netherlands). The time latency to reach the platform was measured (Rzem et al., 2015, Boucherie et al., 2018).

The “Y maze” test was performed to assess short-term memory (Rzem et al., 2015, Lepannetier et al., 2018). The Y maze was made of three identical opaque arms. The test mouse was freely accessible to only two arms for 10 min. After a 30 min inter-trail interval, the third arm was opened and the mouse was put back into the maze. The mouse was video tracked and the time spent in the novel arm was assessed.

“Self-grooming” and “marble burying” tests were performed to assess repetitive behavior. For self-grooming assessment, mice were individually placed in a clear plastic cage for 20min. The first 10 min served as a habituation period and during the second 10 min of testing, time spent grooming were recorded. For the marble burying test, test cages were filled with wood-chip bedding and lightly tamped down to make a flat surface. Twenty glass marbles were placed on the surface in a regular pattern and mice were then placed in the cage. After 30 min, the number of bedding-buried marbles was counted (Amodeo et al., 2012).

### *In utero* electroporation (IUE)

CD1 pregnant females were used for IUE. e13.5 embryos were electroporated with 1 μg/µl of scrambled sh-RNA or sh-Spag5, and 0.5 ug/µl of CAG-GFP in 10 mM Tris buffer (pH 8.0) and 0.01% fast green as described in (Jossin and Cooper, 2011). Needles for injection were pulled from Wiretrol II glass capillaries (Drummond Scientific). Forceps-type electrodes (Nepagene) with 5 mm pads were used for electroporation using the ECM830 electroporation system (Harvard Apparatus). Embryos were collected and fixed at E15.5. sh-RNA were purchased from Origene: scrambled shRNA (Origene, TR30012) and pool of sh-Spag5 (Origene, TR509034) with sequences: TAGTCTCTGGAGACCTGTTGTCCTTGCTT, GGAGGAAGCAATAGAAACAGTGGATGACT, GACAAGTATCTGAGCCATAGGCACATCCT, GCAACAAGGAGCAGGCTACTCAATGGCAA.

### *In-situ* hybridization

RNase free coronal cryosections from e11.5 *Diaph3* KO and control embryos were hybridized with Spag5 fast red-labelled RNAscope probe (Cat No. 505691) as described by the manufacturer.

## Acknowledgements

We thank Dr. Ulrike Gruneberg (University of Oxford, UK) for the Spag5 plasmid, Dr. Mireille Montcouquiol (INSERM, France) for the GPSM2/PINS antibody, Valérie Bonte, Rachid El Kaddouri, Isabelle Lambermont, and Younes Massaoudi for technical support. This study makes use of data generated by the DECIPHER community. A full list of centres who contributed to the generation of the data is available from http://decipher.sanger.ac.uk. Funding for the DECIPHER project was provided by the Wellcome Trust. This work was supported by the following grants: FNRS PDR T00075.15, FNRS PDR T0236.20, FNRS-FWO EOS 30913351, Fondation Médicale Reine Elisabeth, and Fondation JED-Belgique. E.O. L. was supported by “Move-In Louvain” postdoctoral fellowship funded by Marie Skłodowska-Curie Actions of European Commission and Université Catholique de Louvain. G.C. and F.T. are Research Fellow and Research Director at the Belgian Fund for Scientific Research (FNRS).

## Author Contributions

E.O. L. and F.T. designed research and drafted the manuscript.

E.O.L., D. D., Y. J., G. C., O. S., N. T. and P. G. performed the experiments.

All authors edited the manuscript and agreed on its content.

## Competing Interests

Authors declare they have no conflict of interest.

## Supplementary figures and 2 tables

**Figure S1:**
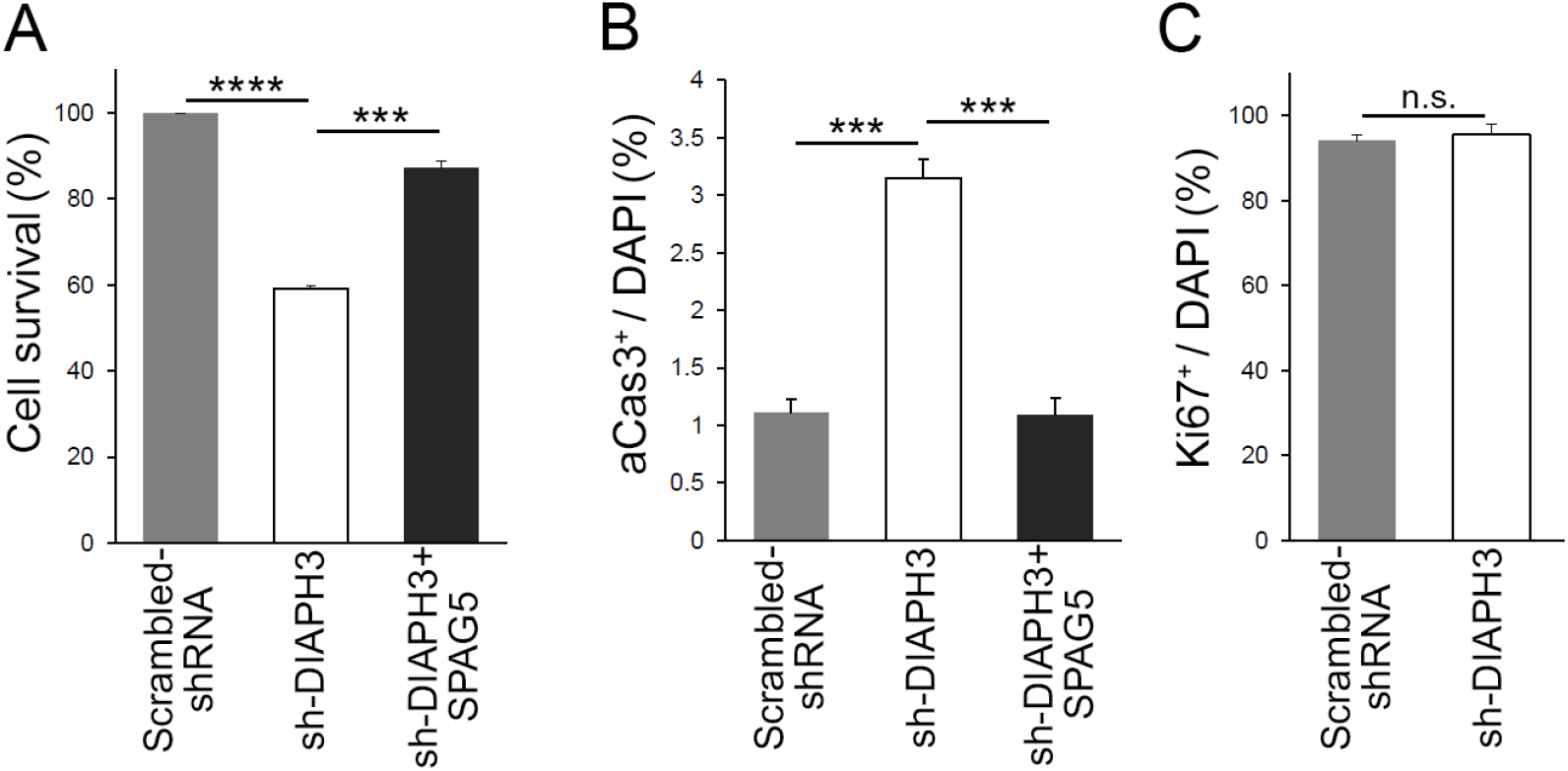
DIAPH3 knockdown reduced cell survival and overexpression of SPAG5 rescued the phenotype. (**A**) sh-DIAPH3 knockdown reduced cell survival to 59.1±0.73% of control cells (transfected with scrambled shRNA, *P*=6.0×10^−7^). Overexpression of SPAG5 rescued cell survival to 87.1±1.7% of control cells (transfected with sh-DIAPH3 and “SPAG5” empty vector, *P*=0.00011). (**B**) Evaluation of cell death by aCas3 immunostaining (3.2±0.16% of cells were apoptotic in sh-DIAPH3 transfections versus 1.1±0.12% in scrambled sh-RNA (*P=*0.00052). Overexpression of SPAG5 restored the percentage of apoptotic cells to normal levels (1.1±0.15% of cells, *P*=0.00071 when compared to sh-DIAPH3 plus empty vector). (**C**) Evaluation of proliferation by Ki67 immunoreactivity (95.5±2.5% after transfection with sh-DIAPH3 versus 94.1±1.5% in cells transfected with scrambled sh-RNA (*P=*0.661). Counts were from 3 independent experiments, Student’s t-test, error bars represent s.e.m.

**Figure S2:**
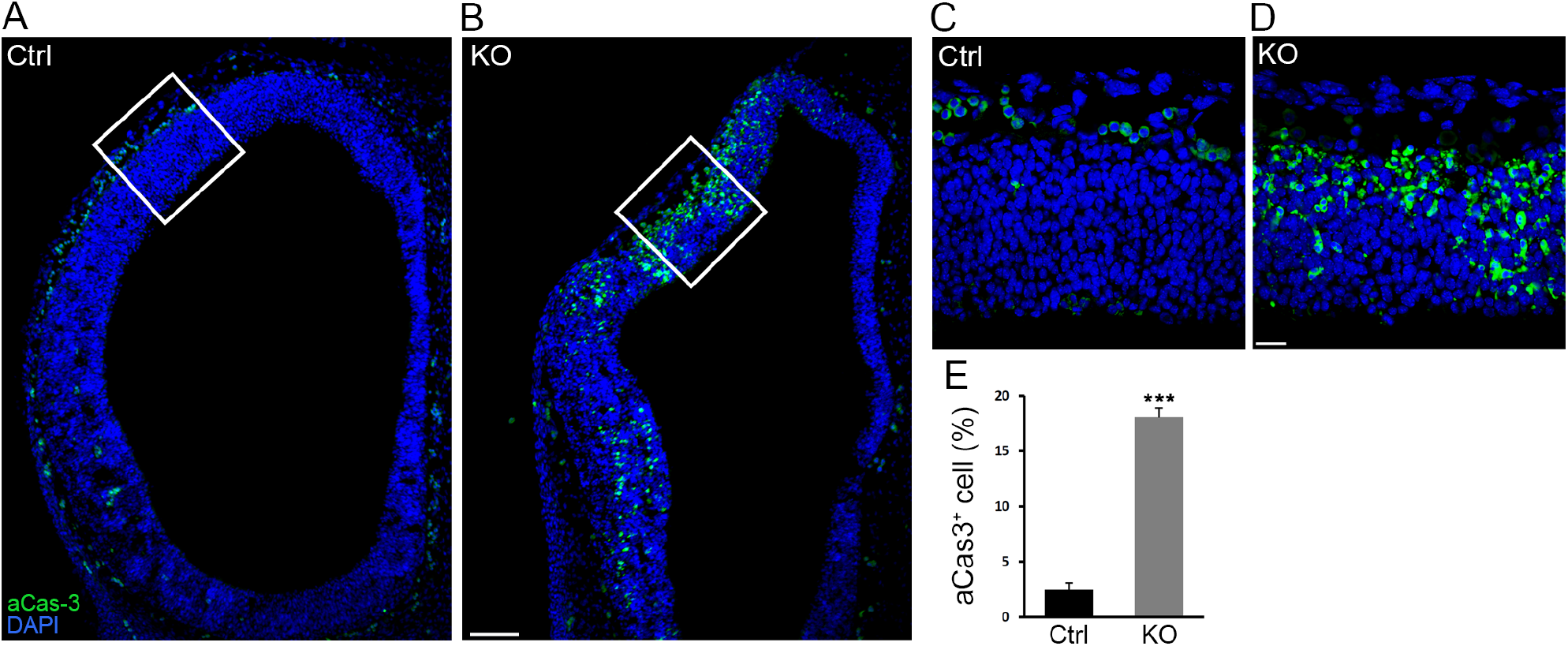
Cell apoptosis in *Diaph3* KO mice. (**A**, **B**) representative images of aCas3 immunostaining (green) on telencephalic sections at e11.5 from control and KO mice illustrating cell death. (**C**) and (**D**) are enlarged images from (**A**) and (**B**) respectively. (**E**) Quantification of apoptotic cells. 18±0.83% of cells in *Diaph3* KO underwent apoptosis versus 2.4±0.65% in control mice. Student’s t-test, *P*=0.00012. Error bars represent s.e.m. Scale bar, 100 µm in (**A**) and (**B**), 20 µm in (**C**) and (**D**). n=9 embryos from 6 different litters

**Figure S3:**
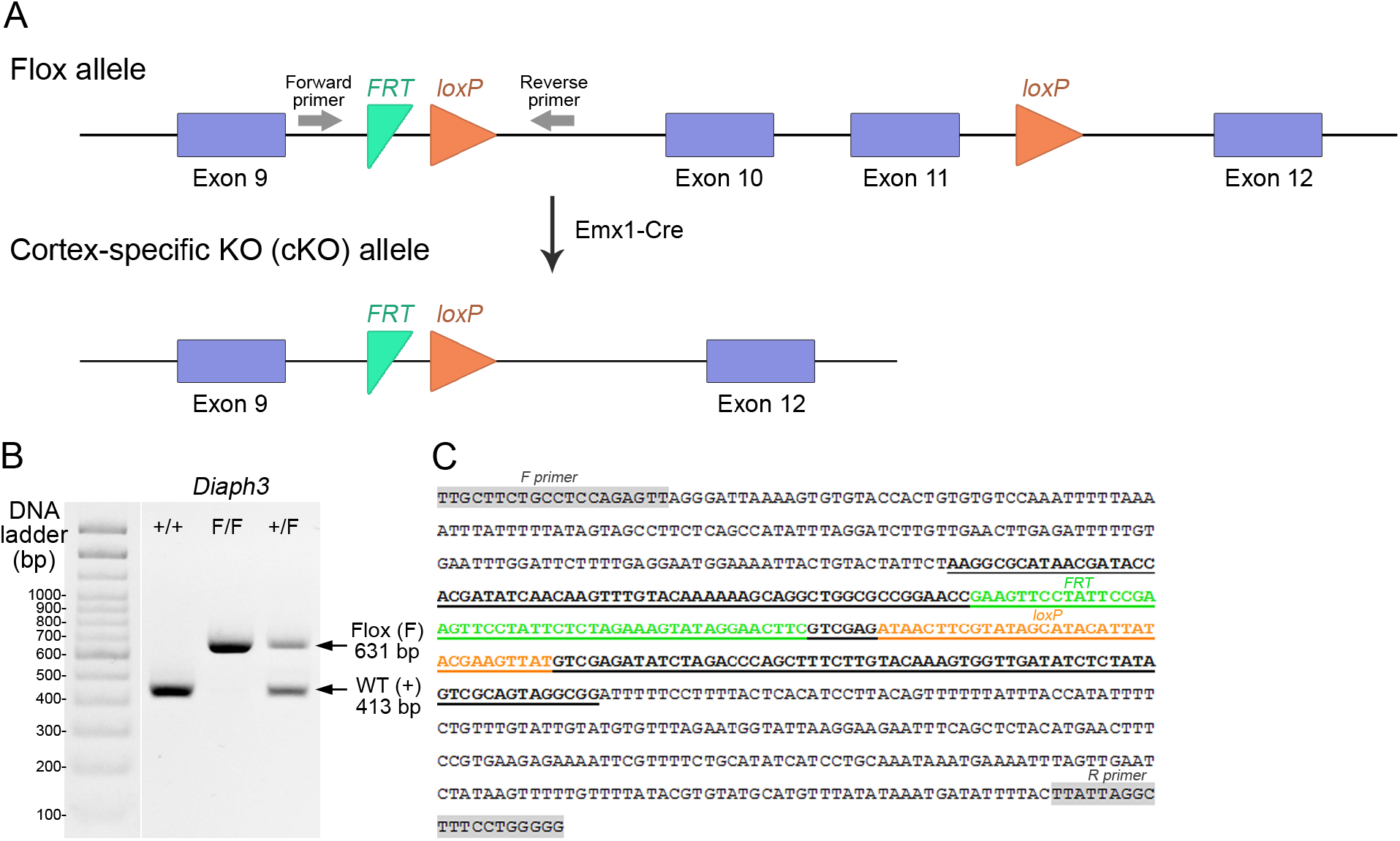
Generation of cortex-specific *Diaph3* KO (cKO) mice. (**A**) Schematic representation of the *Diaph3* conditional allele. Two LoxP sites (Orange arrowheads) were inserted in introns 9 and 11. Exons 10 and 11 were excised in the dorsal telencephalon by crosses with Emx1-Cre mice. (**B**, **C**) Genotyping of *Diaph3 cKO* mice. Forward and reverse primers, flanking the FRT-LoxP cassette in intron 9, amplified a fragment of 413 bp from the wildtype allele (+), and 631 bp from the *Floxed* (*F*) allele (**B**). (**C**) DNA sequence of the recombinant amplicon depicting the FRT (green), LoxP (orange), and PCR primers (grey boxes).

**Figure S4:**
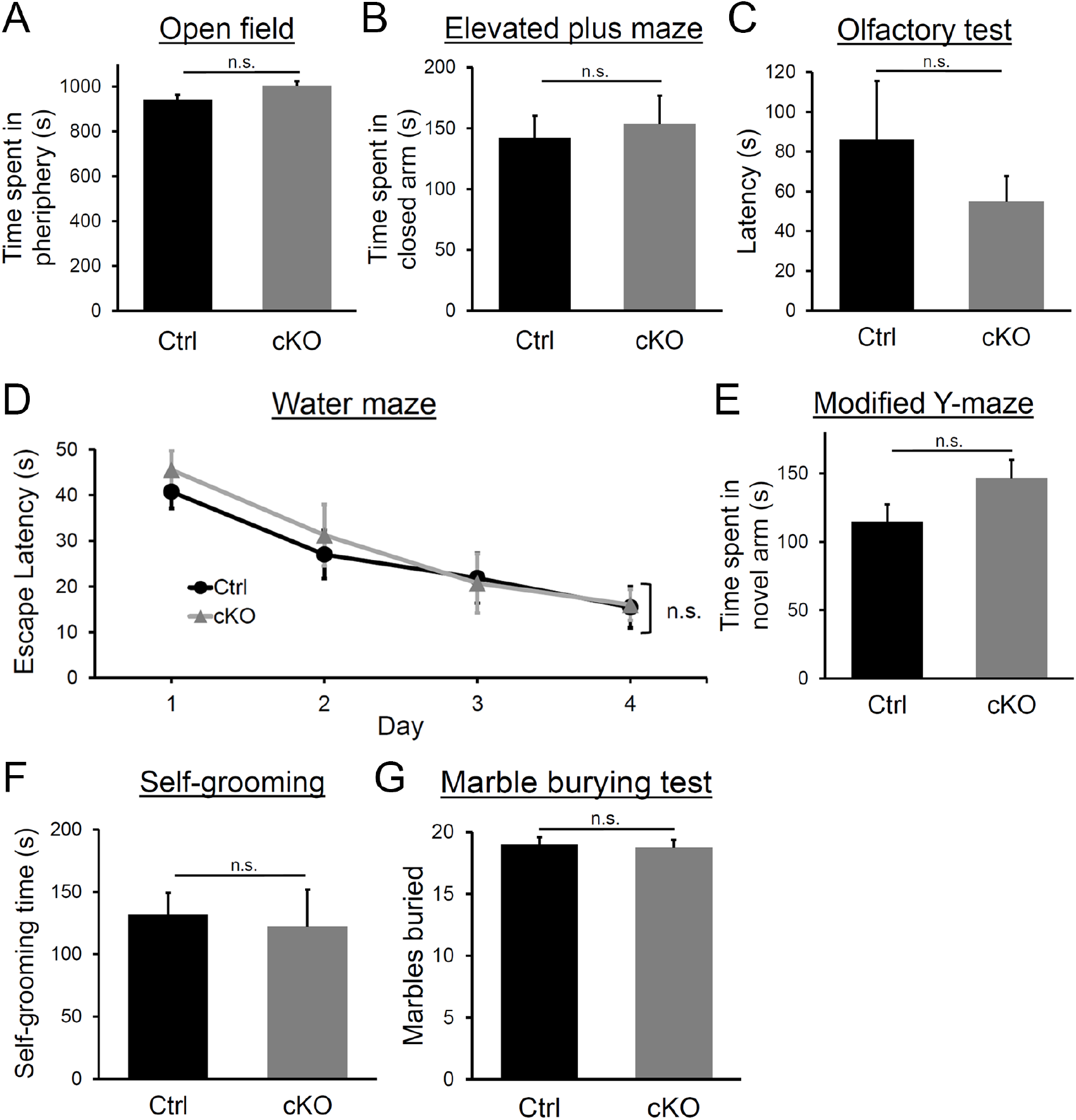
Behavior assessment of *Diaph3* cKO mice. (**A**) Time spent in the periphery of the open field. Student’s *t* test, *P*=0.14. (**B**) Time spent in the closed arm of the elevated plus maze. Student’s *t* test, *P*=0.75. (**C**) Time spent to find the buried pellet in olfactory function test. Student’s *t* test, *P*=0.37. (**D**) Escape latency in the Morris water maze test on the training day (day 1) and three consecutive days (day 2-4). Two-way ANOVA, *P*=0.57. (**E**) Time spent in novel arm in modified Y-maze test. Student’s *t* test, *P*=0.14. (**F**) Time spent for self-grooming. Student’s *t* test, *P*=0.79. (**G**) Number of marbles buried. Student’s *t* test, *P*=0.78, *n* =10 per genotype. Error bars represent s.e.m.

**Figure S5:**
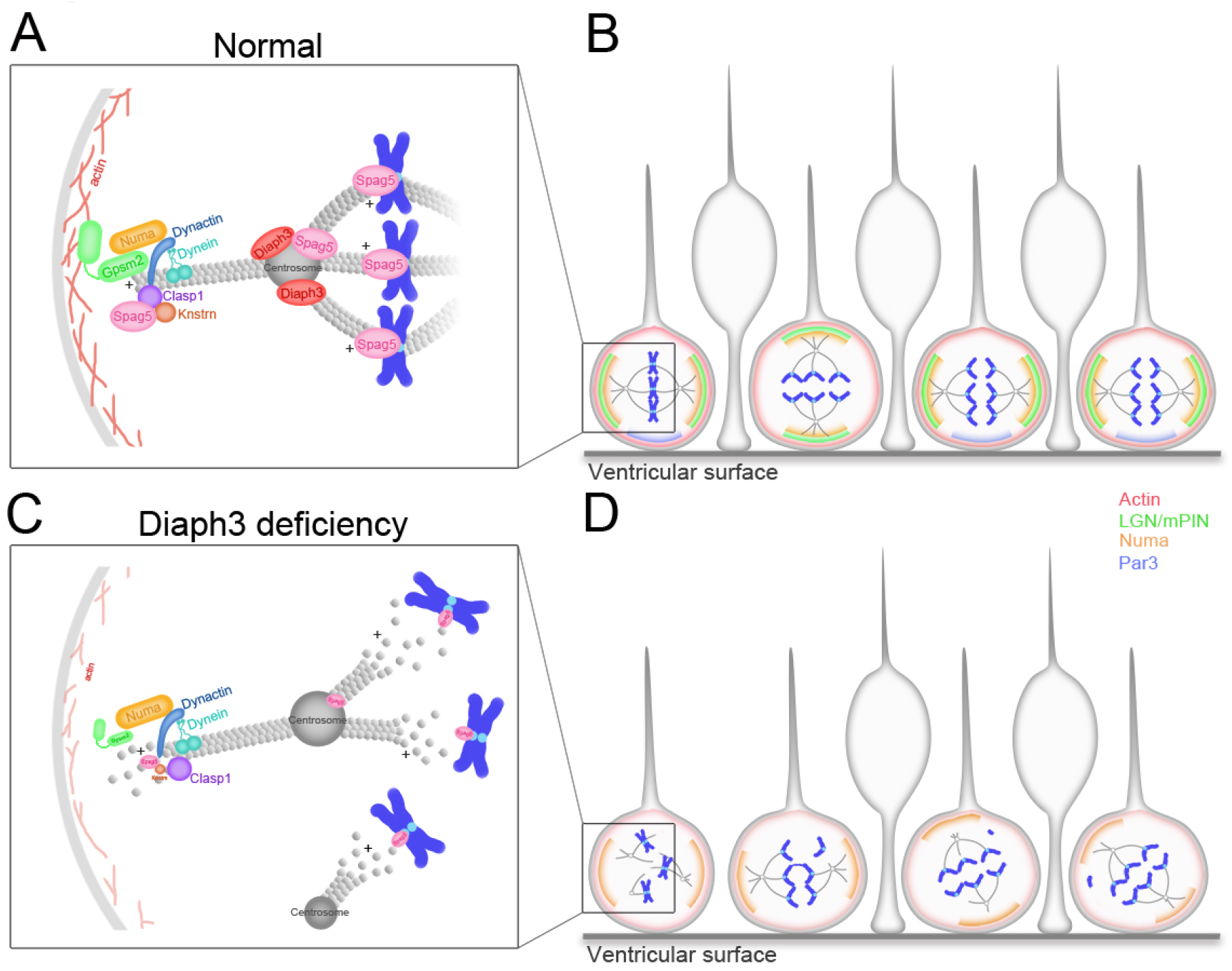
Working model of DIAPH3 function in aNPC. (**A**) During mitosis, DIAPH3 maintains the dynamics of cytoskeleton. Polarity proteins NUMA and GPSM2 assemble underneath cortical F-actin and recruit the dynein/dynactin motor protein complex. Dynein and dynactin interact with SPAG5/KNSTRN/CLASP1 located at the microtubule plus-end and attach astral microtubules to cell cortex, thus providing the pulling force for chromosome bipolar segregation (Okumura et al., 2018, Dunsch et al., 2011, Kern et al., 2016). (**B**) The position of polarity proteins NUMA/GPSM2 directs the orientation of the mitotic spindle and determines the type of division (proliferative versus neurogenic) of aNPC. (**C**, **D**) Absence of DIAPH3 destabilizes actin and microtubules and disrupts the expression of SPAG5, KNSTRN, GPSM2, PAR3 and NUMB, therefore weakening astral microtubules-cell cortex and spindle microtubules-kinetochore interactions. This causes spindle abnormalities, chromosome mis-alignment and mis-segregation, and alters fate decision of aNPC.

**Table S1:** Cell division abnormalities in DIAPH3-knockdown cells. Over 81% of DIAPH3-knockdown cells exhibited mitotic errors (4.2% in control cells). These errors were divided into five “non-exclusive” categories: (1) defect in chromosome alignment (57.4%); (2) abnormal astral microtubules (52.2%); (3) abnormal number of centrosomes (40%); (4) abnormal mitotic spindle (53%); and (5) mis-localization of SPAG5 (60%). *n*=118 cells transfected with scrambled-shRNA, and 115 with sh-DIAPH3 from 5 individual experiments.

|  | Scrambled-shRNA (%) | sh-DIAPH3<br>(%) |
| --- | --- | --- |
| Abnormal cell division | 4.2 | 81.7 |
| (1) defect in chromosome alignment: lagging<br>chromosomes and/or asymmetric metaphase plate | 4.2 | 57.4 |
| (2) abnormal astral microtubule | 0.0 | 52.2 |
| (3) abnormal number of centrosomes | 0.0 | 40.0 |
| (4) abnormal mitotic spindle | 0.0 | 53.0 |
| (5) mis-localization of SPAG5 | 0.0 | 60.0 |

**Table S2:**
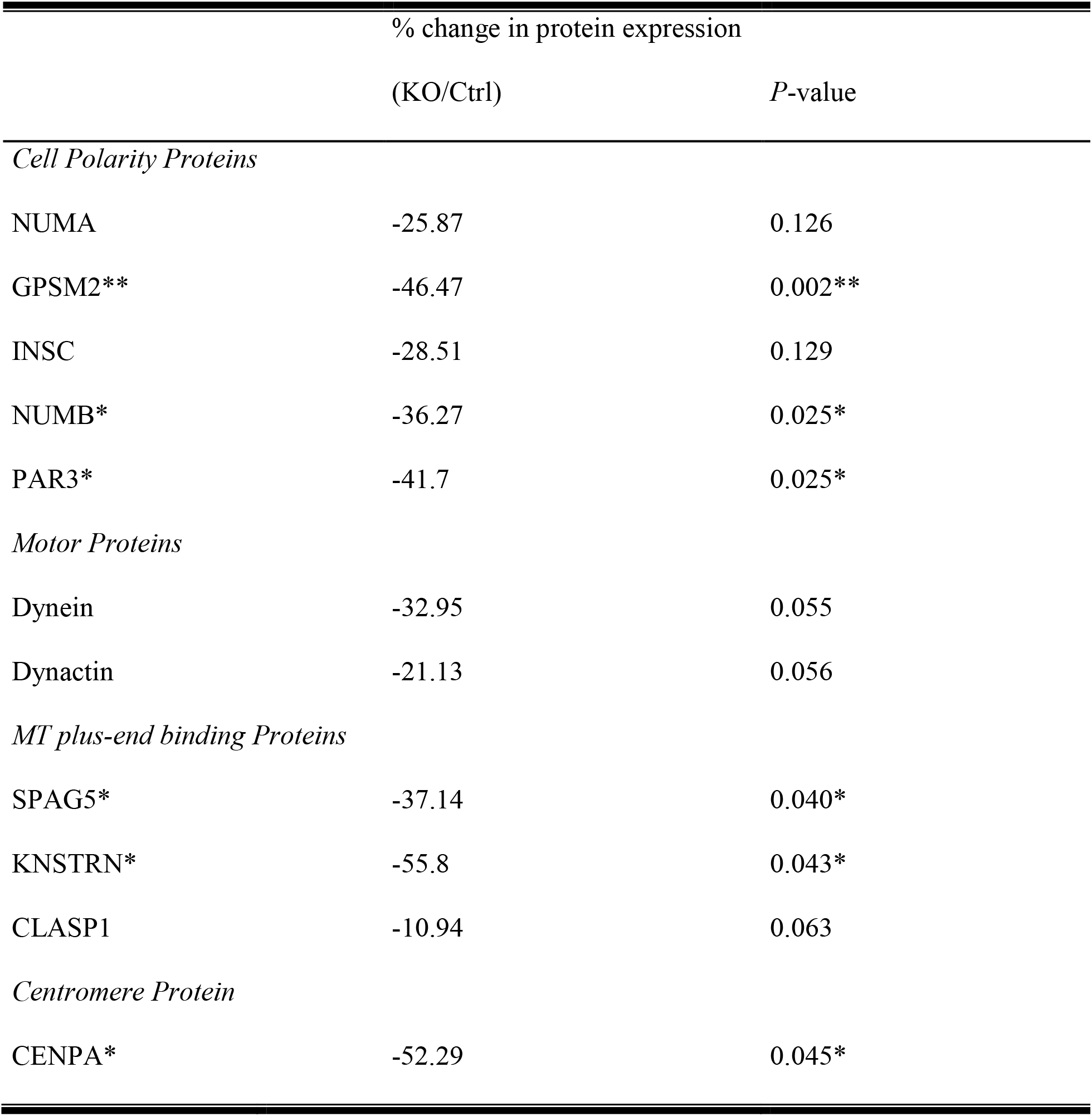
Expression of proteins related to spindle orientation in *Diaph3* KO and control mice. *n* =4 embryos for each genotype (3 different litters), Student’s *t*-test, \**P*<0.05, ** *P*<0.01.

## Notes

### Competing Interest Statement

The authors have declared no competing interest.

